# Comprehensive analysis of genomic diversity of SARS-CoV-2 in different geographic regions of India: An endeavour to classify Indian SARS-CoV-2 strains on the basis of co-existing mutations

**DOI:** 10.1101/2020.07.14.203463

**Authors:** Rakesh Sarkar, Suvrotoa Mitra, Pritam Chandra, Priyanka Saha, Anindita Banerjee, Shanta Dutta, Mamta Chawla-Sarkar

## Abstract

Accumulation of mutations within the genome is the primary driving force for viral evolution within an endemic setting. This inherent feature often leads to altered virulence, infectivity and transmissibility as well as antigenic shift to escape host immunity, which might compromise the efficacy of vaccines and antiviral drugs. Therefore, we aimed at genome-wide analyses of circulating SARS-CoV-2 viruses for the emergence of novel co-existing mutations and trace their spatial distribution within India. Comprehensive analysis of whole genome sequences of 441 Indian SARS-CoV-2 strains revealed the occurrence of 33 different mutations, 21 being distinctive to India. Emergence of novel mutations were observed in S glycoprotein (7/33), NSP3 (6/33), RdRp/NSP12 (4/33), NSP2 (2/33) and N (2/33). Non-synonymous mutations were found to be 3.4 times more prevalent than synonymous mutations. We classified the Indian isolates into 22 groups based on the co-existing mutations. Phylogenetic analyses revealed that representative strain of each group divided themselves into various sub-clades within their respective clades, based on the presence of unique co-existing mutations. India was dominated by A2a clade (55.60%) followed by A3 (37.38%) and B (7%), but exhibited heterogeneous distribution among various geographical regions. The A2a clade mostly predominated in East India, Western India and Central India, whereas A3 clade prevailed in South and North India. In conclusion, this study highlights the divergent evolution of SARS-CoV-2 strains and co-circulation of multiple clades in India. Monitoring of the emerging mutations would pave ways for vaccine formulation and designing of antiviral drugs.

## 1. Introduction

When a virus adapts to a new host within an endemic setting, it needs to exploit the host’s cellular machinery for successful entry, establishing its replication and evading host’s immune responses [1]. To achieve this, viruses try to modify antigenic epitopes on virus encoded proteins by continuously mutating its genome. As the virus evolves in a stable environment with minimum selection process, transition mutations are more frequent than the transversions [2]. Deleterious mutations which may hamper virus’s life cycle are filtered out through natural selection pressure. The mutations which confer some advantage to the virus persist and evolve further. Thus, digging deep into the nature of mutations can decipher how selection pressure might be acting on this novel virus.

RNA viruses display a characteristic feature of high mutability and with no exception, SARS-CoV-2 being a positive strand RNA virus has been evolving at a rapid rate since its emergence in Wuhan during the end of 2019. In the span of six months (December, 2019-June, 2020), the circulating SARS-CoV-2 strains have accumulated a large number of mutations which might result in altered virulence, infectivity and transmissibility [3, 4]. Evolutionary behaviour of viruses frequently relies on co-occurrence of multiple mutations in different genes or within a single gene. Continuous monitoring of these single nucleotide polymorphisms and locating them to the protein coding genes might help to gain insight into the genetic diversity of SARS-CoV-2. Accumulation of mutations in other respiratory viruses, like Influenza have shown to result in generation of vaccine escape mutants or drug resistant mutants leading to continuous need of developing new vaccines or drugs [5]. In the context of urgent requirement of effective vaccine or antiviral drug against SARS-CoV2, it is imperative to monitor the evolving mutations in the viral proteins.

Hence, this study was designed to analyse and compare the genetic mutations among SARS-CoV-2 viruses across various geographical regions of India against the prototype ‘Wuhan strain’ [6]. Establishing an atlas of co-existing mutations across SARS-CoV-2 genome might underscore their genetic evolution within the various epidemiological settings of India.

## 2. Materials and Methods

### 2.1. Sequence retrieval

Full genome nucleotide sequences of 441 SARS-CoV-2viruses circulating in India (Mar-May 2020) were retrieved from the GISAID repository [7] (Supplementary Table 1). Several other clade-specific reference gene sequences of SARS-CoV-2 were also downloaded from GISAID for construction of the dendrogram.

### 2.2. Screening of mutations and phylogenetic analyses

The novel mutations within the Indian SARS-CoV-2 isolates were analysed with respect to the prototype strain “Wuhan-Hu-1” (MN908947.3). The phylogenetic dendrogram was constructed based on the whole genome of 22 representative Indian strains and 10 reference sequences, using MEGA-version X (Molecular Evolutionary Genetics Analysis), recruiting the maximum-likelihood statistical method at 500 bootstrap replicates, using the best fit nucleotide substitution model (General Time Reversible). MUSCLE v3.8.31 was used for multiple sequence alignment. Amino acid sequences were retrieved through TRANSEQ (Transeq Nucleotide to Protein Sequence Conversion Tool, EMBL-EBI, Cambridgeshire, UK).

## 3. Results

### 3.1. Identification and analyses of various mutations among the SARS-CoV-2 strains circulating in different geographical regions of India

To unravel the mutations accumulating through natural selection across SARS-CoV-2 genome, we performed a meticulous whole-genome sequence analysis encompassing 441 Indian SARS-CoV-2 strains deposited in the GISAID repository (Supplementary Figure 1). A total of 33 different mutations existed among the 441 Indian isolates in comparison to the prototype strain Wuhan-Hu-1 (mutations those were found in minimum 5 isolates were only considered). The S protein harboured 8 substitution mutations, of which 6 were non-synonymous (G21724T/L54F, A21792T/K77M, G21795T/R78M, G23311T/E583D, A23403G/D614G, G23593T/Q677H) and 2 were synonymous (C22444T/D294D, C23929T/Y789Y). 7 mutations were found across NSP3 protein: 6 non-synonymous (G4866T/G716I, C4965T/T749I, C5700A/A994D, A6081G/D1121G, C6310A/S1197R, C6312A/T1198K) and 1 synonymous mutation (C3037T/F106F). The RdRp/ NSP12, NSP2, N and NSP4 proteins have acquired 5 (C13730T/A97V, C14408T/P323L, C14425A/L329I, G15451A/G571S, G16078A/V880I), 4 (C884T/R27C, G1397A/V198I, C1707T/S301F, G1820A/G339S), 3 (T28311C/P13L, C28854T/S194L, GGG28881AAC/RG203KR) and 2 mutations (G8653T/M33I, C8782T/S76S), respectively. Single mutations have been identified in the 5’-UTR region (C243T), NSP6 (G11083T/L37F), ORF3a (G25563T/Q57H) and ORF8 (T28144C/L84S) sequences. Rest of the genome was found to be conserved, having no significant amino acid substitutions. S, NSP3 and NSP12 proteins are found to be more susceptible to mutations followed by NSP2, N, NSP4, NSP6, ORF3a and ORF8 (Figure 1A). Four mutations-C241T in 5’-UTR (n=242/441), C3037T/F106F in NSP3 (n=241/441), C14408T/P323L in RdRP (n=240/441), and A23403G/D614G in S (n=239/441)-were found to predominated mainly in East, West and Central India (Figure 1B). Subsequent leading mutations were G11083T/L37F in NSP6 (n=163/441), C13730T/A97V in RdRP (n=160/441), C23929T/Y789Y in S (n=152/441), T28311C/P13L in N (n=158/441) and C6312A/T1198K in NSP3 (n=144/441), which prevailed mostly across South and North India (Figure 1B-G). G25563T/Q57H in ORF3a (n=91/441) was the next preponderant mutation in India, principally Western India, followed by G21724T/L54F and C22444T/D294D in S (n=54/441 and 40/441, respectively), C28854T/S194L in N (n=39/441), C6310A/S1197R in NSP3 (n=33/441), T28144C/L84S in ORF8 (n=30/441), C8782T/S76S in NSP4 (n=30/441) and GGG28881AAC/RG203KR in N (n=28/441) (Figure 1B, D). Maximum amino acid variations were observed in NSP3, NSP4, RdRp, S and N (Figure 1D) genes of strains circulating in Western India, whereas North India exhibited highest mutations in NSP2 (Figure 1F).

**Figure 1.**
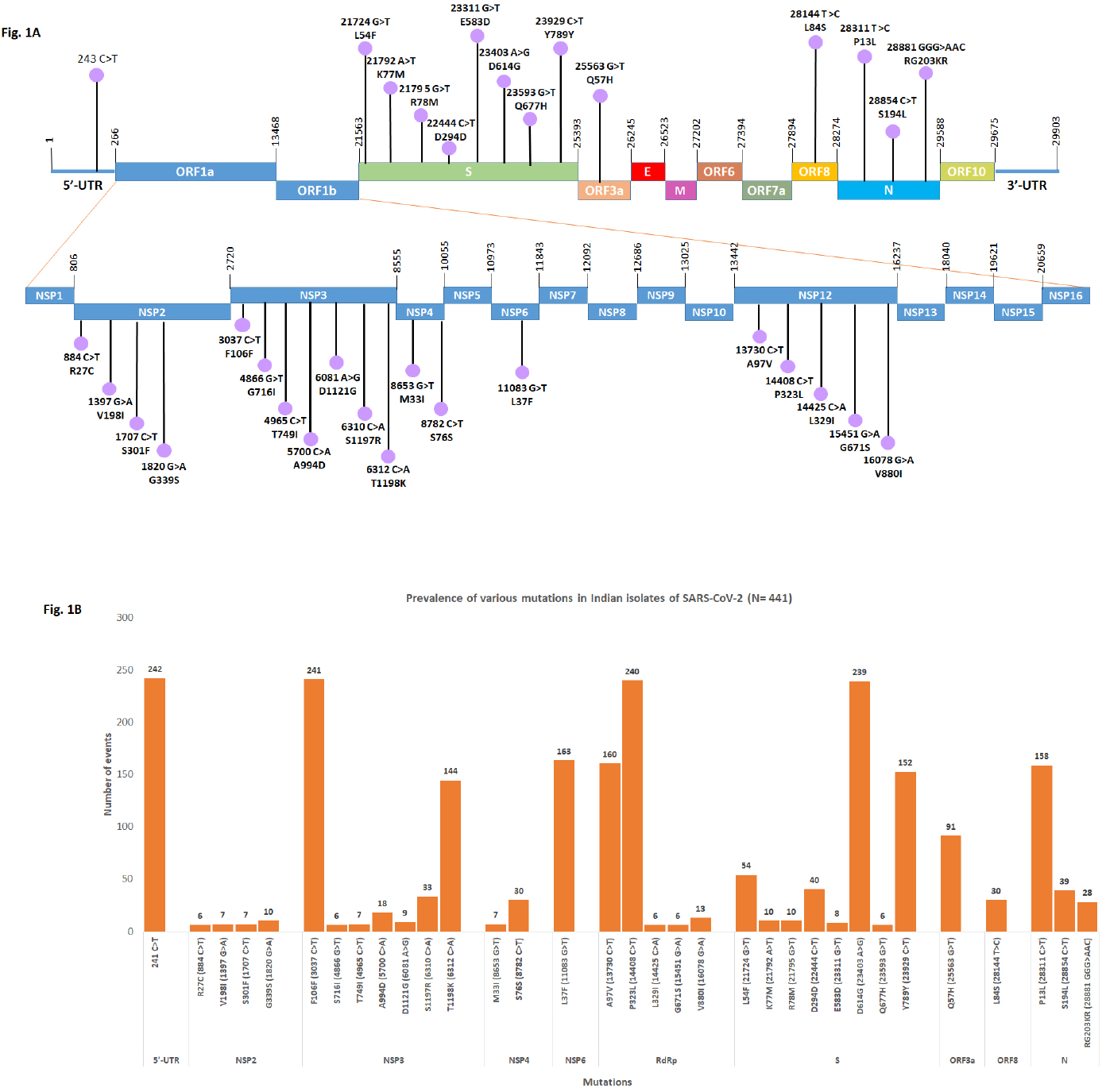
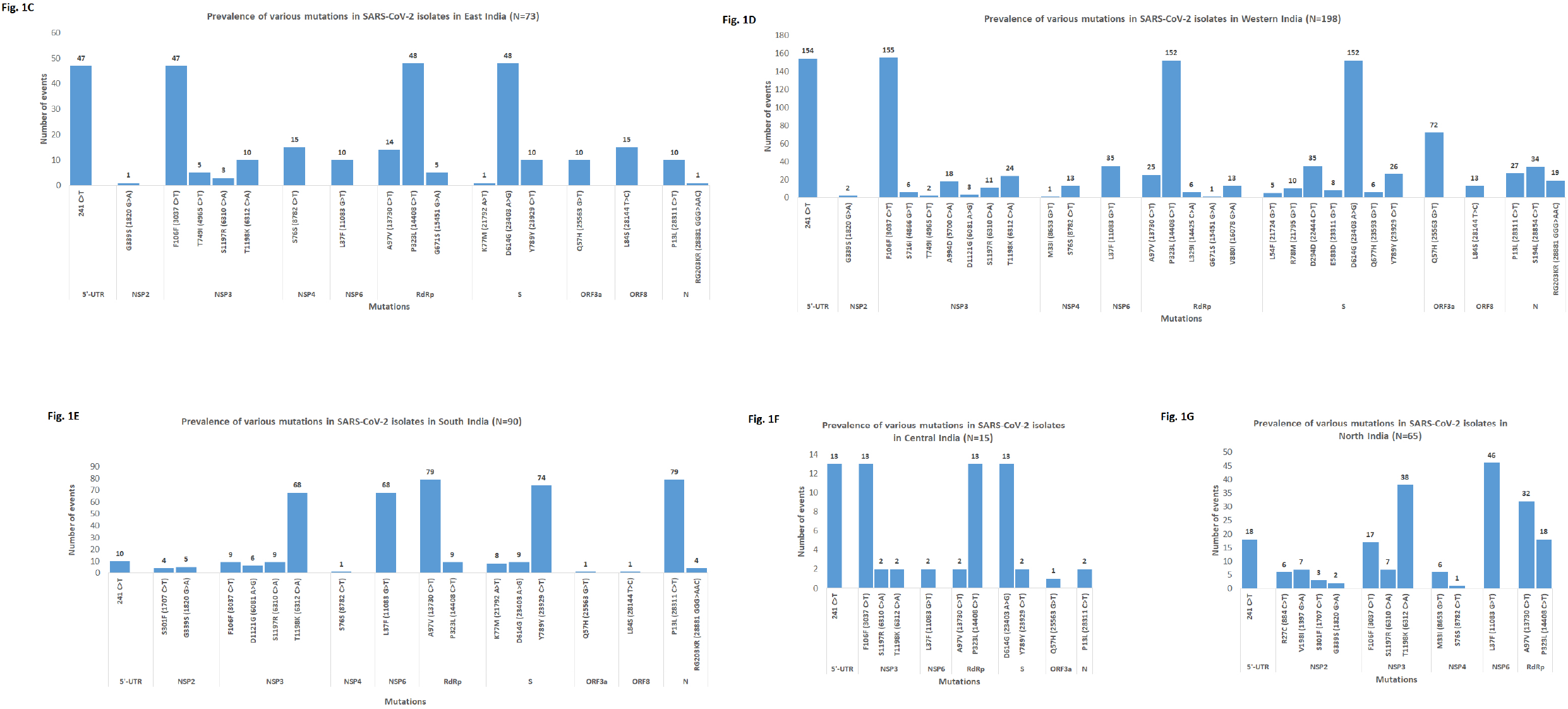
**(A-B):** Identification of various mutations present in the genome of SARS-CoV-2 circulating in India. (A) Pictorial presentation of 33 different mutations (represented at the level of nucleotide as well as amino acid) found in different regions (coding and non-coding regions) of SARS-CoV-2 genome. (B) Relative frequencies of 33 different mutations in India. **(C-G):** Identification of various mutations present in the genome of SARS-CoV-2 circulating in different geographic regions in India. Relative frequencies of various mutations in (C) East India, (D) Western India, (E) South India, (F) Central India and (G) North India.

### 3.2. Emergence of synonymous and non-synonymous mutations: Analysis of nucleotide substitution events (transition and transversion) at the level of codon positions

Analysis of mutational events per sample revealed that maximum number of Indian isolates harboured 5 mutations followed by 4, 6, 7 and 2 mutations (Figure 2A). Non-synonymous mutations occurred 3.24 times (1504/463) more frequently than synonymous mutations (Figure 2B). We have identified 8 nucleotide substitutions: 4 transitions (C>T, A>G, G>A, T>C) and 4 transversions (G>T, C>A, G>C, A>T) which are responsible for 29 non-synonymous and 4 synonymous mutations (Figure 2C). C>T transition was the most prevalent substitution (Figure 2C), occurring predominantly in the second position of the codon followed by the third position (Figure 2D). On account of C>T transition, 6 non-synonymous mutations (A97V, P13L, S194L, S301F, T749I) ensued in the second position of codon, whereas 4 synonymous mutations (D294D, Y789Y, F106F, S76S) occurred in the third position (Figure 2D). The G>T transversion was the next dominant nucleotide substitution (occurring frequently in the third position and rarely in the second position of the codon) which generated 6 (E583D, Q667H, L54F, M33I, L37F, Q57H) and 2 (R78M, S716I) non-synonymous mutations. The third most prevalent A>G transition occurred solely at the second position of the codon and was responsible for D614G and D1121G mutations. The C>A transversion, was seen more frequently in the second position of the codon generating T1198K and A994D mutations, though it did appear occasionally in the first position causing L329I and S1197R mutations. G>T transversion exclusively occurred in the first position of the codon (V880I, G671S, V198I and G339S mutations). The co-frequent nucleotide substitutions, T>C (2nd position) and G>C (1st position of codon) generated the L84S and G204R mutations. G>A transversion (both in the 2nd and 3rd positions) fostered the R203K change (Figure 2C-D).

**Figure 2:**
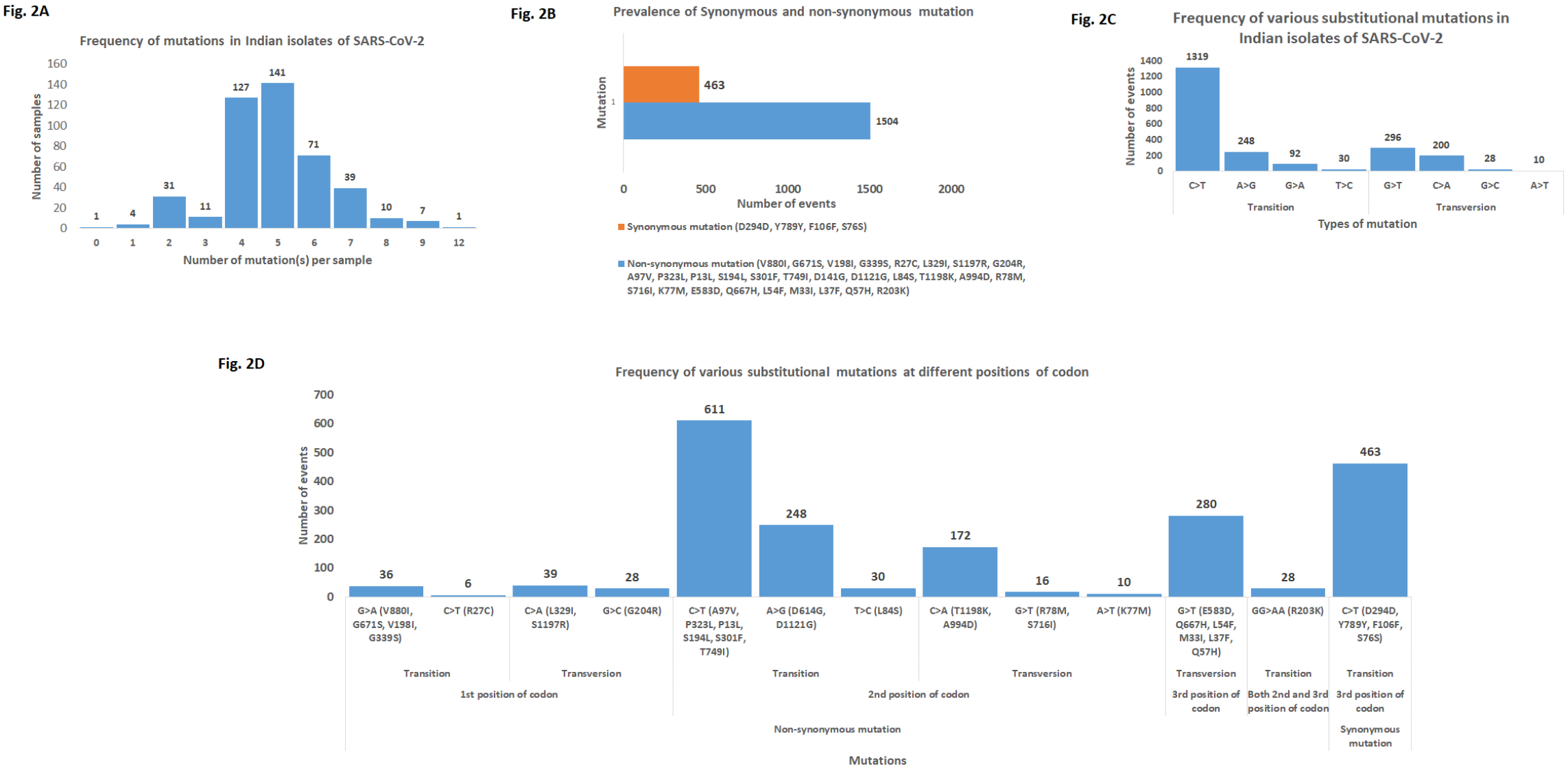
Analysis of synonymous and non-synonymous mutations regarding nucleotide substitutions at different positions of codon. (A) Frequency distribution of SARS-CoV-2 isolates harbouring varying numbers of co-existing mutations. (B) Prevalence of synonymous and non-synonymous mutations in SARS-CoV-2 genomes across India. (C) Frequency distribution of various transitional (C>T, A>G, G>A and T>C) and transversional (G>T, C>A, G>C and A>T) substitution events. (D) Frequency distribution of various types of substitutional events occurred at different nucleotide positions (1^st^, 2^nd^ and 3^rd^) of the codon.

### 3.3. Spatial classification of the Indian SARS-CoV-2 strains based on co-existing mutations and their preponderance in different geographical regions across India

On the basis of co-existing mutations, we could classify the 428 Indian isolates into 22 groups, each group representing a different set of co-existing mutations (Figure 3A, Table 1). Out of 22 groups, 12 groups represented the strains belonging to the A2a clade (most prevalent) having the clade specific 4 mutations (D614G/S, F106F/NSP3, C241T/5’-UTR and P323L/RdRp). 11 out of 12 groups have acquired additional mutations (Q57H/ORF3a, S194L/N, D294D/S, V880I/RdRp, E583D/RdRp, L54F/S, R78M/S, RG203KR/N, A994D/NSP3, G671S/RdRp, A97V/RdRp, L291I/RdRp) in various combination. The group having only four characteristic mutations is the most predominant one among A2a clade and the leading group in India (Figure 3A). The groups with four mutations along with Q57H or Q57H, S194L and D294D or RG203KR and A994D are moderately dominant among A2a clade. Eight groups represented the A3 clade bearing L37F mutation along with various combinations of V198I/NSP2, M33I/NSP4, R27C/NSP2, P13L/Y789Y/S, A97V/RdRp, T1198K/NSP3, S1197R/NSP3, S301F/NSP2, G339S/NSP2, D1121G/NSP3 and K77M/S. The group having the co-existing mutations like L37F, P13L, Y789Y, A97V and T1198K is the second most dominant group among A3 clade and second leading group in India. Two groups represented the strains of B clade having the characteristic L84S/ORF8 and S76S/NSP4 with or without T749I/NSP3. Overall, India is dominated by A2a (55.60%) followed by A3 (37.38%) and B (7%) clades (Figure 3B). Geographical distribution revealed the predominance of A2a clade strains across East, West and Central India; whereas A3 clade were common in South and North India. The B clade strains have been exclusively reported from East and West India (Figure 4A-F).

**Figure 3:**
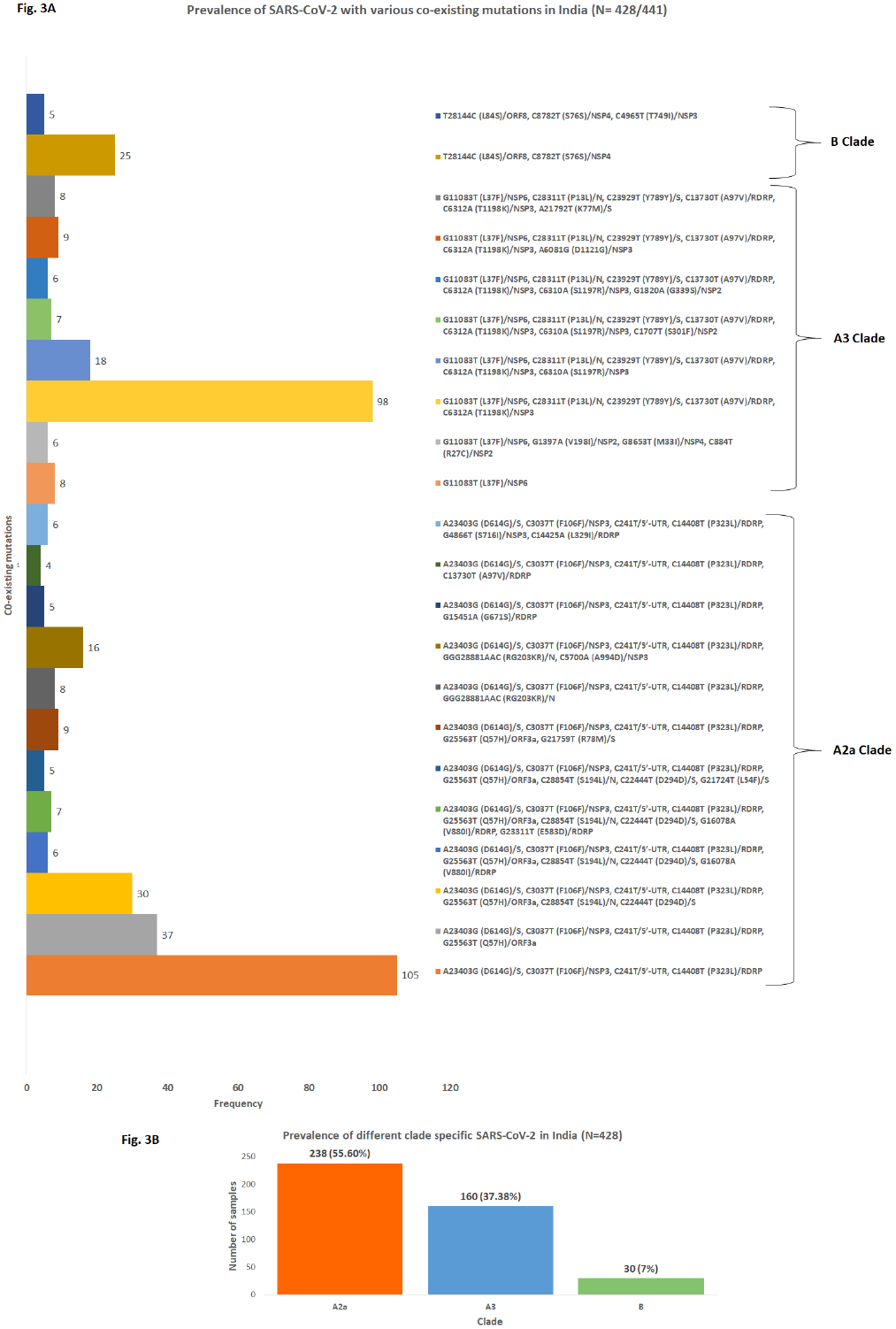
Grouping of SARS-CoV-2 strains on the basis of co-existing mutations and analysis of their prevalence. (A) Mutational analysis revealed the presence of the three clade (A2a, A3 and B) specific SARS-CoV-2 strains in India. Accumulation of novel mutations in addition to clade specific changes directed us to classify A2a clade strains into 12 groups, A3 clade strains into 8 groups and B clade strains into 2 groups. We also shown the number of strains belonging to each group. (B) Prevalence of three clade specific mutations in India. A2a clade (55.60%) was found to be most prevalent in India, followed by A3 (37.38%) and B (7%).

**Figure 4.**
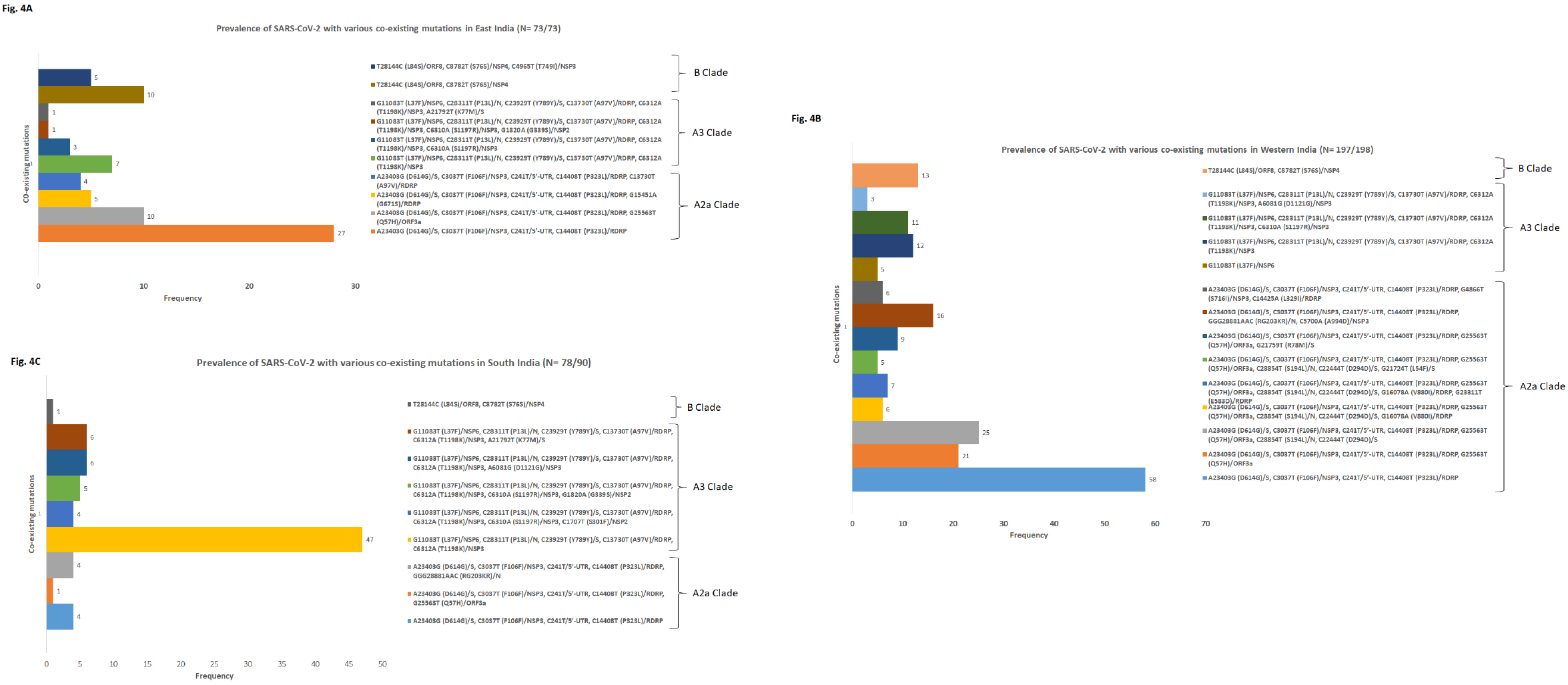
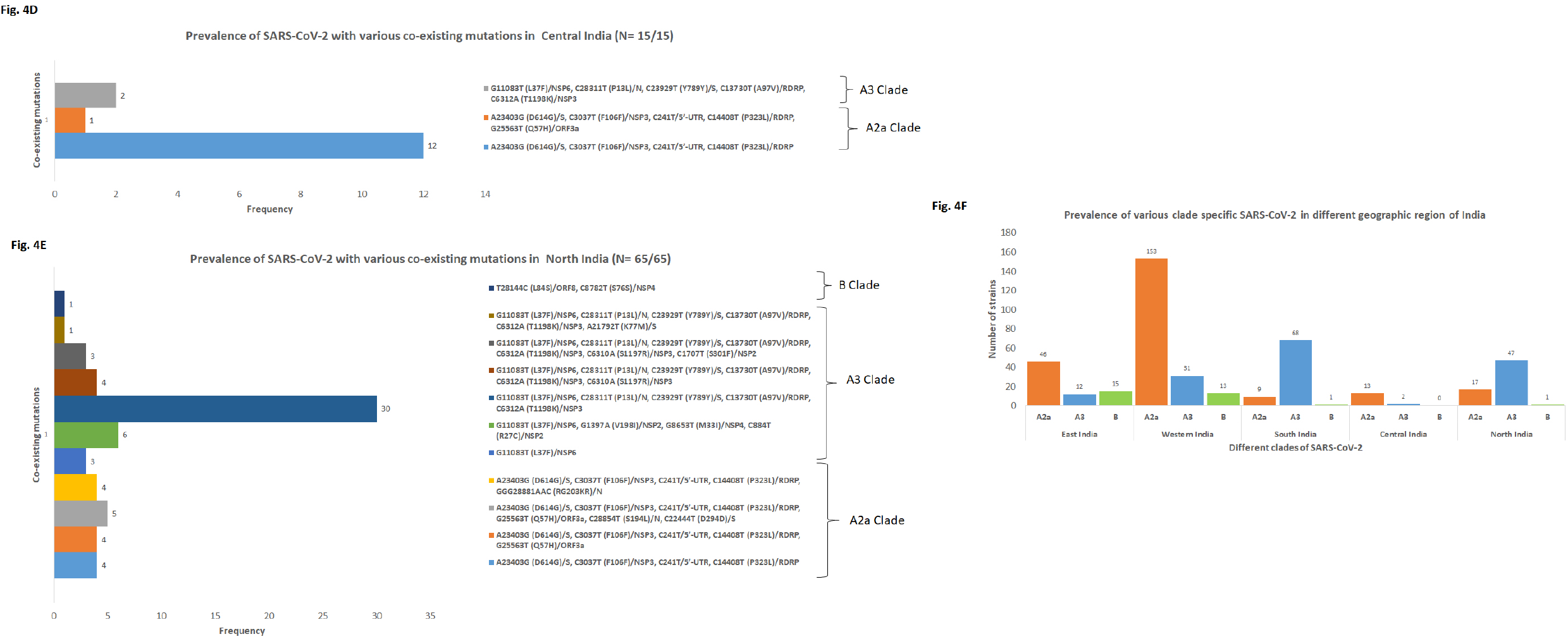
**(A-C):** Prevalence of three different clades (A2a, A3 and B) and their sub groups in different geographic regions in India. Frequency distribution of strains belonging to each groups of three different clades in (A) East India, (B) Western India and (C) South India, **(D-F):** Prevalence of three different clades (A2a, A3 and B) and their sub groups in different geographic regions in India. Frequency distribution of strains belonging to each groups of three different clades in (D) Central India and (E) North India. (F) Prevalence of three different clades in different geographic regions of India.

**Table 1.**
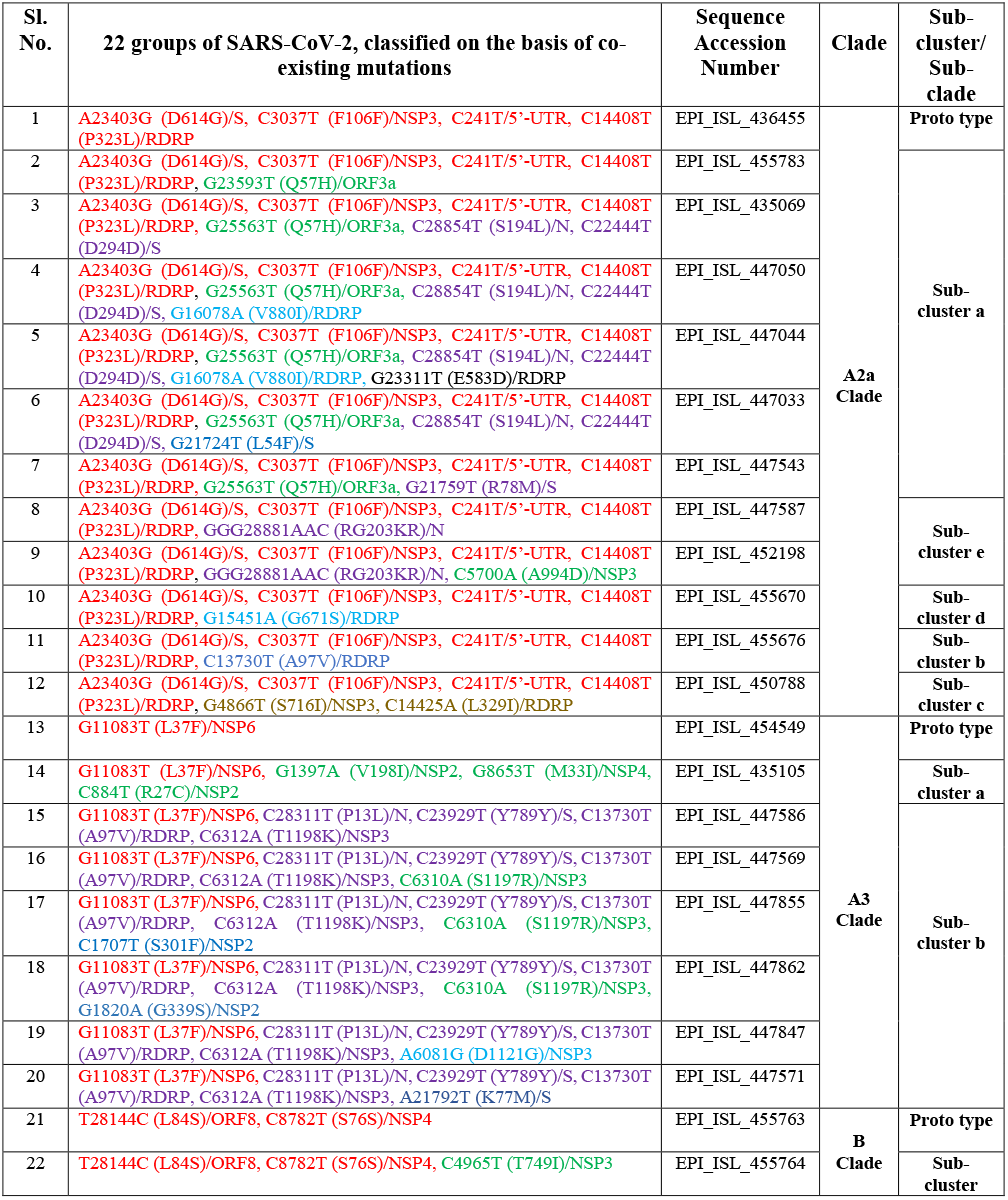
List of accession numbers of representative strains of 22 groups of SARS-CoV-2.

### 3.4. Phylogenetic analysis of 22 groups of Indian SARS-CoV-2 isolates in comparison to the various clade specific strains

The genetic closeness and the clustering pattern of the 22 groups of Indian isolates was analyzed, by comparing with the various SARS-CoV-2 clade-specific strains and the prototype clade O strain from Wuhan (MN908947.3). Whole genome sequences of 22 representative Indian strains (one from each of the 22 co-evolving mutant groups) (Table 1) along with strains denoting 10 different clades were selected for phylogenetic analyses. As expected, the dendrogram revealed that the 22 isolates clustered with strains of 3 different clades (12 strains with A2a, followed by 8 with A3 and only 2 with B4-2 clade). The prototype strain (clade O) belonged to the lineage harbouring A3 and B clade strains. Very interestingly, the 22 Indian strains, representing different groups, have generated sub-clusters within their respective clades, based on the accumulations of co-existing mutations in addition to the clade-specific mutations (mentioned on the branches of each lineage) (Figure 5, Table 1). Within the A3 clade, 2 sub-clusters were seen: a (1 strain) and b (6 strains), bearing 3 (V198I, M33I, R27C) and minimum 4 (P13L, Y789Y/S, A97V, T1198K) coexisting mutations, respectively in addition to the characteristic L37F mutation. Within clade-B4-2, only 1 representative strain with T749I mutation in addition to clade-specific L84S and S76S mutations was observed. Indian strains within clade A2a formed 5 sub-clusters (viz, a-6 strains; b, c, d-1 strain each and e with 2 strains). In addition to the A2a clade-specific 4 mutations (D614G, F106F, C241T P323L); other novel variations like: 1 (Q57H), 1 (A97V), 2 (S716I and l329I), 1 (G671S) and 2 (RG203KR) were revealed in sub-cluster a-e strains, respectively. All the representative Indian strains had >99% nucleotide sequence homology among themselves. The prototype strain belonging to the O clade clustered close to the A3 clade (>98% identity).

**Figure 5:**
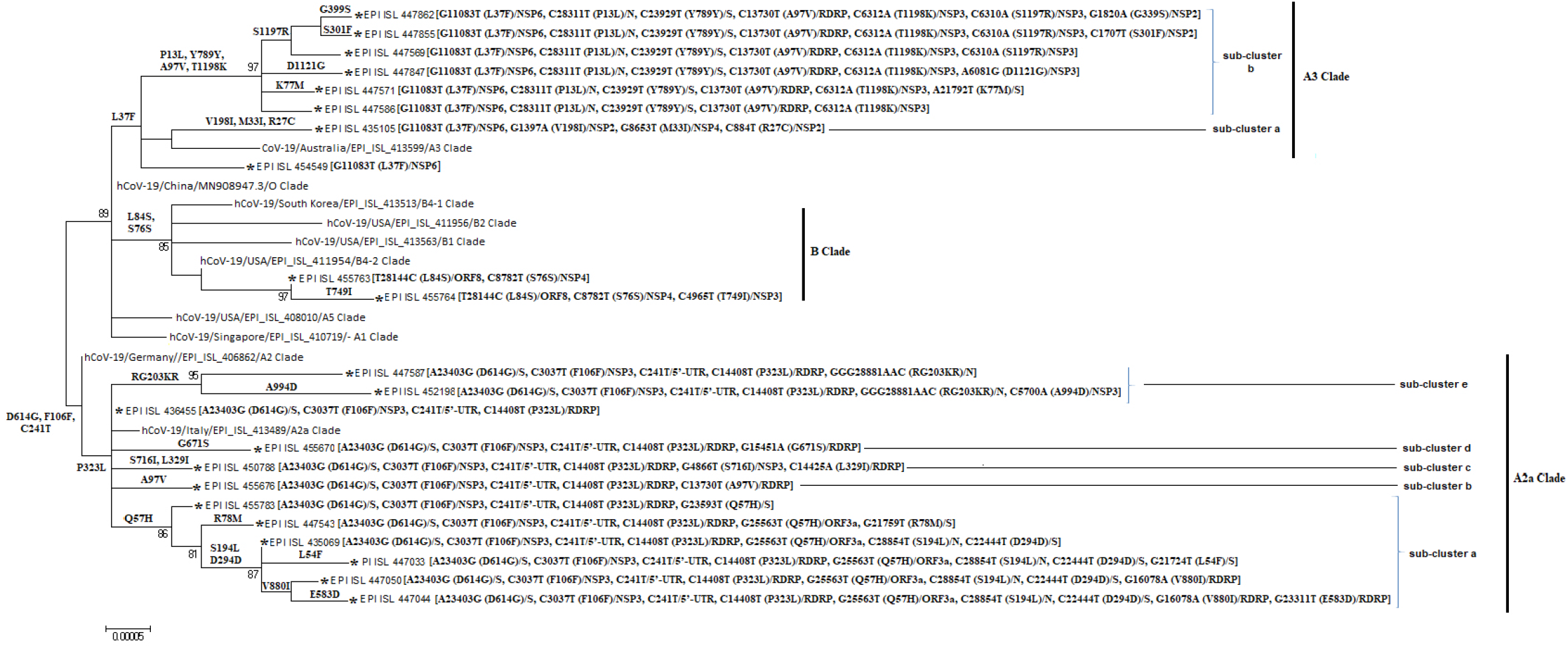
Molecular phylogenetic analysis by maximum likelihood method. Phylogenetic dendrogram based on whole genome sequences of 22 representative strains from 22 different groups along with 9 clade specific known strains and the prototype O-clade strain (MN908947.3). 22 representative strains have been marked with star (*). Scale bar was set at 0.00005 nucleotide substitution per site. Bootstrap values of less than 70% are not shown. The best fit model used for constructing the phylogenetic dendrogram was General Time Reversible Model (GTR).

## 4. Discussion

During the emergence of SARS-CoV-2 virus in Wuhan, a monophyletic-clade O prevailed. As the virus spread across the continents, it started accumulating mutations to adapt in various epidemiological settings. In the present study, we performed a comprehensive mutational analysis of 441 Indian SARS-CoV-2 strains identified in different geographical regions of India, and classified them on the basis of co-existing mutations.

Our data highlighted the existence of 33 different mutations (32 mutations in 9 different protein coding genes and 1 in 5’-UTR) among Indian SARS-CoV-2 strains. Maximum number of mutations were detected in S protein (8) followed by NSP3 (7), NSP12 (5), NSP2 (4), N (3) and NSP4 (2). Only single mutation has been observed in 5’-UTR, NSP6, ORF3a and ORF8 gene segments. Along with 28 non-synonymous mutations, we observed four silent mutations (D294D/S, F106F/NSP3, S76S/NSP4 and Y789Y/S) which may not have any apparent effect on protein structure, but may amend codon usage and have repercussion on translation efficiency [8]. Mutation in the 5’-UTR region may have significant impact on folding, transcription and replication of viral genome. Comparing the mutation patterns of India and abroad, we witnessed certain mutations in S (L54F, K77M, R78M, D294D, E583D, Q677H, Y789Y), NSP3 (G716I, T749I, A994D, D1121G, S1197R, T1198K), RdRP (A97V, L329I, G571S, V880I), NSP2 (S301F, G339S) and N (P13L, S194L) which are unique to Indian isolates. D614G/S, a characteristic mutation of A2 clade, has been found to correlate strongly with the increased case fatality rate [9]. Recent reports also suggested that D614 residue remains embedded in an immuno-dominant linear epitope of S protein and displayed exaggerated serological response. The D614G mutation has also been established to be associated with reduced sensitivity of neutralizing antibodies toward S protein [10, 11]. Among the 5 novel non-synonymous mutations in S protein, L54F, K77M and R88M were found to reside within the NTD domain of S1 subunit and may have significant effect on the receptor binding ability of S1 subunit [12]. Two mutations E583D and Q677H were observed in the linker region of S1 and S2 subunits and may influence host protease mediated cleavage of S1 and S2 subunit during entry of the SARS-CoV-2 [12]. Genomic integrity of SARS-CoV-2 principally relies on the functional efficiency of the RdRp/NSP12. We observed the presence of A97V and L329I changes in the NiRaN domain, V880I in the thumb domain and G571S in the finger domain of RdRP, which could compromise its replication-fidelity and also alter its sensitivity towards inhibitors like Remdesivir, Ribavirin and Favipiravir which are recommended for COVID-19 treatment [13]. The S194L mutation resides in the central region of N protein which is essential for its oligomerization [14, 15]. We observed the codominance of 4 mutations (C24lT/5’-UTR, D614G/S, F106F/NSP3 and P323L/RdRp) in India, being most prevalent in the East, West and Central India. This is followed by a group of 5 co-dominating mutations (L37F/NSP6, T1198K/NSP3, A97V/RdRp, Y789Y/S and P13L/N), having higher frequency in South and North India.

Traditionally, rapidly mutating positive-sense single stranded RNA viruses harbour a higher transitional load in their genomes than transversions [16]. Consistent with this report, we also noticed 3.24 times more frequent transitional events over transversion in SARS-CoV-2 genome. Out of 33, 14 mutations were found to be derived from transversions and rest of the 19 mutations originated due to transitions. As transversion events radically change the properties (size/charge/polarity) of the substituted amino acid, any such mutation in the coding region could change the protein function. It is not surprising to observe 5 transversion mutations in S protein because viral capsid proteins often undergo greater number of mutations leading to altered functions. This might be an immune-elusive viral strategy as maximum neutralizing antibodies are generated as well as vaccines are designed against the surface protein epitopes.

Digging deep into the nature of mutations we found C>T transition was accountable for 12 of the 34 reported mutations (considering RG203KR as R203K and G204R) followed by G>T (8/34), G>A (6/34) and C>A (4/34). Least frequent substitutions were A>G (2/34), A>T (1/34), T>C (1/34) and G>C (1/34). High frequency of C>T and G>A transitions could drive from APOBEC (Apolipoprotein B mRNA Editing Catalytic Polypeptide-like), a family of cytidine deaminase, mediated C to U deamination [17]. Furthermore, maximum C>T changes have been identified at the 2^nd^ position of the codon (harbouring Cs as in GCN [Ala] to GUN [Val], CCN [Pro] to CUN [Leu], UCN [Ser] to UUN [Leu/Phe], CAN [Thr] to AUN [Ile]), further underscoring the role of APOBECs which prefers 5’-NCU-3’ sites for action [18]. All the synonymous mutations resulted from C>T transitions occurring at 3 position of the codon. The A>G mutation (responsible for A2 clade specific D614G mutation) and T>C mutation (responsible for B clade specific L84S mutation) could have arisen as a result of the ADAR (Adenosine Deaminase Acting on RNA) effect. Thus synthetic inhibitors of APOBEC and ADAR might prove better amongst the arsenal of anti-SARS-CoV-2 drugs under trial.

Since the outbreak in Wuhan, the unceasing accumulation of genetic mutations has driven the formation of multiple clades and subclades from the prototype clade-O. Congregation of the two mutations, L84S (T28144C) in ORF8 and S76S (C8782T) in NSP4, led to the emergence of clade-B; which has been circulating more in North and South America but less in countries of Africa and Europe. The world’s leading A2-clade emerged upon accumulation of three mutations-D614G (A23403G) in S, F106F (C3037T) in NSP3, and C241T in 5’-UTR. In contrast to the B clade, A2 clade was the dominant one in Europe, Africa, Asia and Oceania but was less frequent in North and South America. After inclusion of an additional mutation P323L (C14408T) in RdRP of A2 clade, a more preponderant sub-clade A2a was established. A2a clade has most likely originated in Asia, nevertheless, it has rapidly transmitted to Europe and America and has become the cardinal clade. The A3 clade, having the signature L37F (G11083T) mutation in NSP6, is distributed maximally within Singapore, Brunei, Thailand, Indonesia and a few parts of Middle-East including Iraq, Iran and Kyrgyztan [4, 8, 19].

In this study, we have observed the heterogeneous distribution of SARS-CoV-2 strains of three different clades (A2a, A3 and B) in different geographic regions of India. The A2a clade (55.60%), the leading clade in India, is predominant in West, East and Central India and but less frequent in North and South India. The A3 clade (37.38%) is India’s second most prevalent clade, largely prevails in North and South India and is less frequent in East, West and Central India. The B clade (7%), the least frequent one, has been principally reported from East and Western India. We classified the Indian isolates into 22 groups on the basis of co-existing mutations. 12 groups represented the strains of A2a clade having four common characteristics mutations along with various amalgamation of novel mutations mostly associated with ORF3a, RdRp, S and N proteins. 8 groups aligned with A3 clade having L37F characteristic mutation in concert with several unique mutations typically linked to nonstructural proteins (NSP2, NSP3, NSP4 and NSP12). 2 groups represented B clade strains and are found to be associated with the novel mutation T749I/NSP3.

The SARS-CoV-2 genome is accumulating mutations at a very high frequency. As suggested by the ‘mutation-selection balance’ and the ‘speed-fidelity trade-off’ theories [20, 21], this might be because it has concentrated its endeavours to hasten its replication for increased host-transmissibility at the cost of accurate replication. This might be advantageous during adaptation within a heterogeneous population where it is undergoing strong directional selection pressure due to host immunity [22]. However, other factors governing this response might be the viral genomic constellation, presence of RNA secondary structures, influence of host RNA editing enzymes (ADAR and APOBEC) and genetic hitchhiking [23, 24]. Not all mutations are in favour of the virus. It has been reported that when the beneficial mutations surpass the detrimental effects of associated deleterious mutations, then the deleterious mutations are subject to fixation, especially when it is being encoded from the same DNA [25, 26]. In this case, they are synonymous with respect to all the non-structural proteins being encoded from ORF 1a and 1b. Though the strain ‘O’ was the first SARS-CoV-2 strain which was responsible for introduction of SARS-CoV-2 infection in humans, it is being replaced eventually by its swarm of circulating viral quasispecies [27, 28], at the face of host immune pressure where the virus is utilising its fast replication strategy to enhance its propagation.

In conclusion, the present study highlights the rapid accumulation of various novel mutations in several proteins, principally in S glycoprotein and RdRp, that has led to the indigenous convergent evolution of SARS-CoV-2 circulating in different geographic regions of India. Presently, vaccine development and RdRp inhibitors based therapies are being targeted to control this global pandemic situation. However, for successful therapeutics, it would be imperative to monitor the mutations in targeted genes. This study has provided the much needed information regarding the novel mutations in S, RdRp and several other non-structural proteins, which could pave ways for vaccine formulation and for designing of antiviral drugs targeting specific viral proteins.

## Acknowledgement

The study was supported by Indian Council of Medical Research, India. RS is supported by Senior Research Fellowship from UGC. The authors acknowledge The hard work of scientists and laboratory staffs in all the COVID-19 testing laboratories and Next gen sequencing labs across India.

---

**Supplementary Figure 1:**
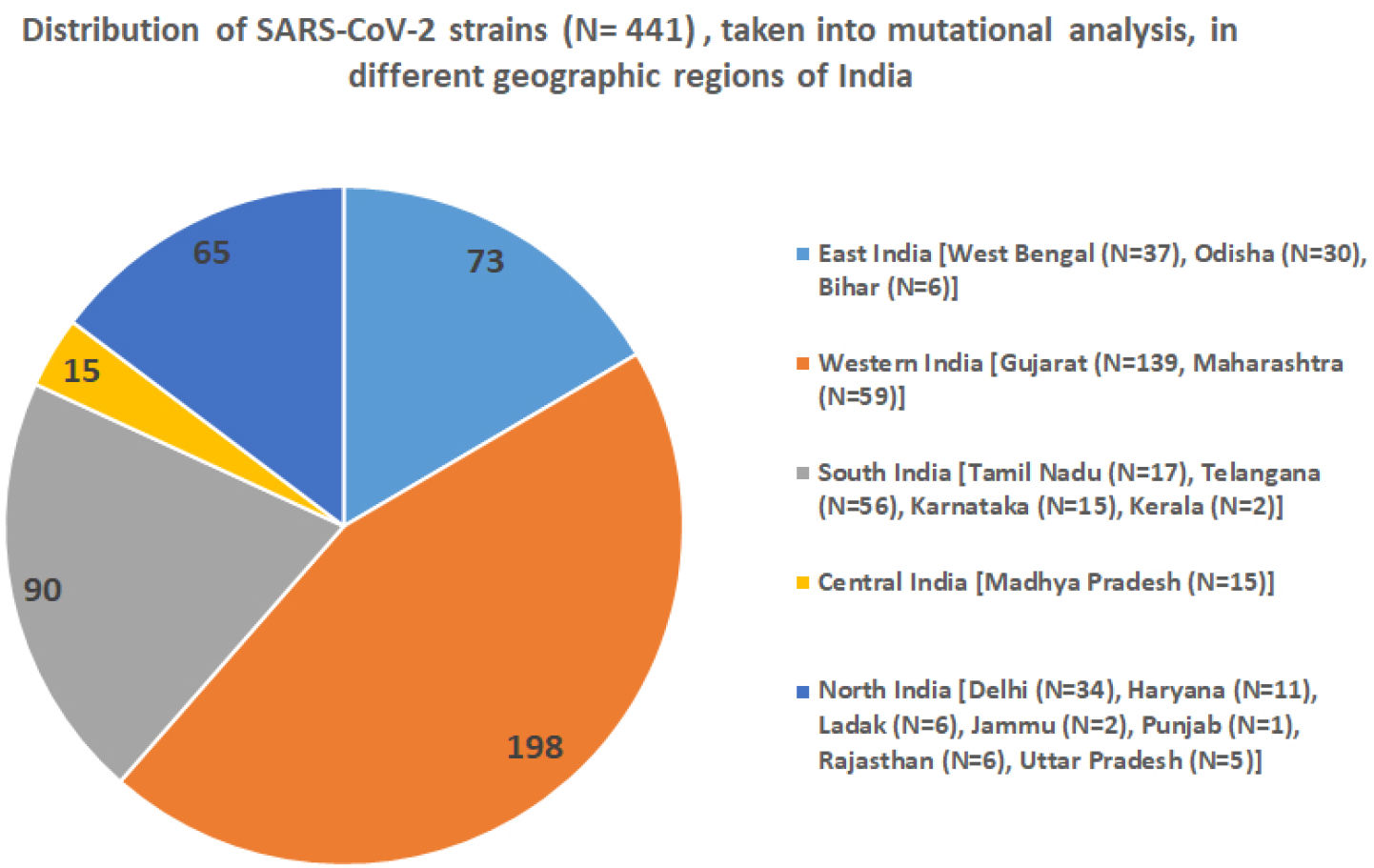
Pie chart representation of strains from different geographic region taken into consideration for mutation analysis. States belong to each such region and numbers of strains taken are mentioned as well.

**Supplementary Table 1:**
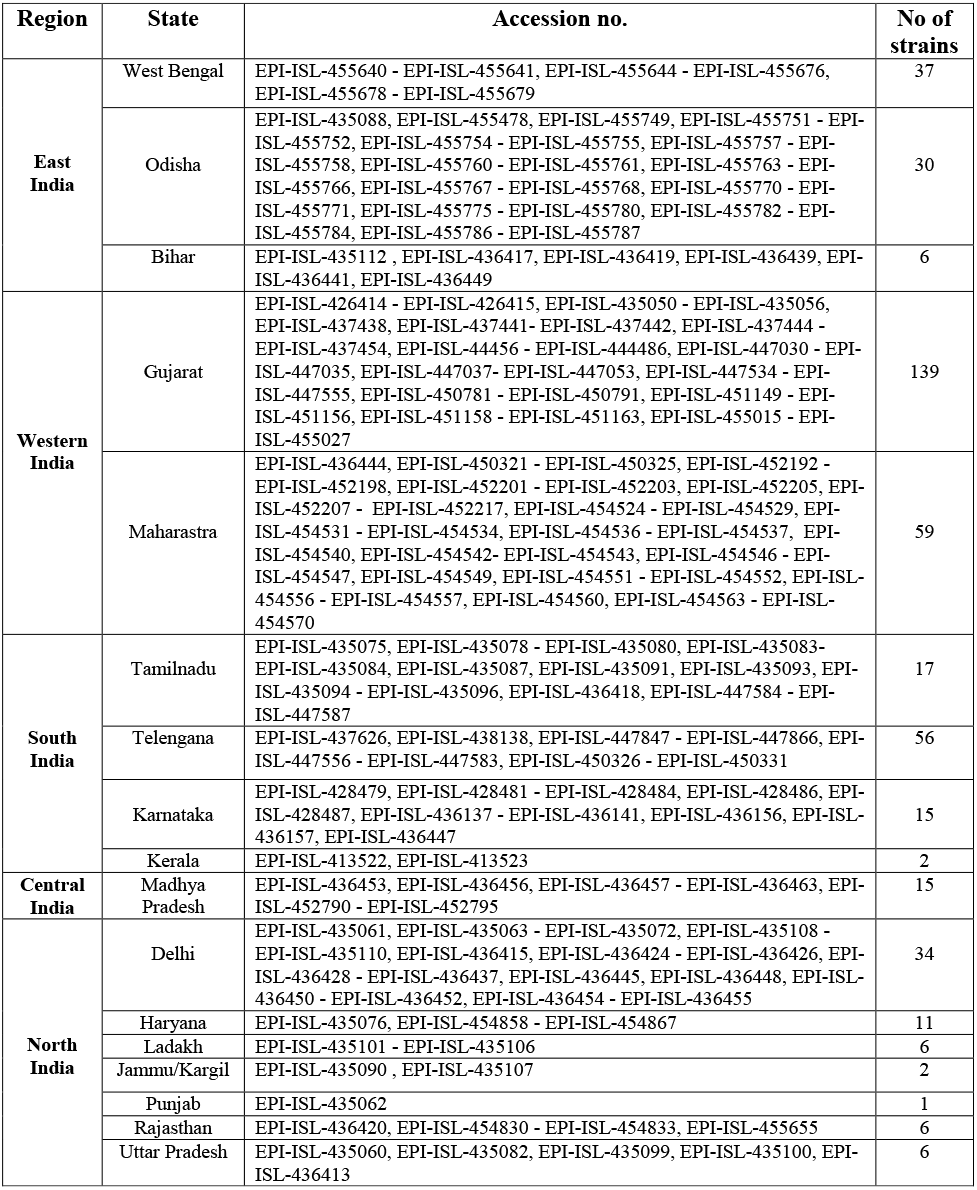
List of accession numbers of 441 SARS-CoV-2 strains taken into mutational analysis.

## References

1. Sackman AM, McGee LW, Morrison AJ, et al. Mutation-Driven Parallel Evolution during Viral Adaptation. Mol Biol Evol. 2017;34(12):3243–3253.

2. Barr, J. N., and R. Fearns. “Genetic Instability of RNA Viruses.” Genome Stability. Academic Press, 2016. 21–35.

3. Maitra A, Sarkar MC, Raheja H, Biswas NK, Chakraborti S, Singh AK, Ghosh S, Sarkar S, Patra S, Mondal RK, Ghosh T. Mutations in SARS-CoV-2 viral RNA identified in Eastern India: Possible implications for the ongoing outbreak in India and impact on viral structure and host susceptibility. J. Biosci. 2020;45(1).

4. Gomez-Carballa A, Bello X, Pardo-Seco J, Martinon-Torres F, Salas A. The impact of super-spreaders in COVID-19: mapping genome variation worldwide. bioRxiv 097410 (Preprints) 2020. [Cited June 30, 2020]. Available from: https://www.biorxiv.org/content/10.1101/2020.05.19.097410v3.full.

5. Du X, Wang Z, Wu A, Song L, Cao Y, Hang H, Jiang T. Networks of genomic cooccurrence capture characteristics of human influenza A (H3N2) evolution. Genome research. 2008 Jan 1;18(1):178–87.

6. Wu D, Wu T, Liu Q, Yang Z. The SARS-CoV-2 outbreak: What we know. Int J Infect Dis. 2020;94:44–48.

7. Shu, Y., McCauley, J. (2017) GISAID: Global initiative on sharing all influenza data - from vision to reality. EuroSurveillance, 22(13).

8. Mercatelli, D.; Giorgi, F.M. Geographic and Genomic Distribution of SARS-CoV-2 Mutations. Preprints 2020, 2020040529. doi: 10.20944/preprints202004.0529.v1.

9. Becerra-Flores M, Cardozo T. SARS-CoV-2 viral spike G614 mutation exhibits higher case fatality rate [published online ahead of print, 2020 May 6]. Int J Clin Pract. 2020;e13525.

10. Korber B, Fischer W, Gnanakaran SG, Yoon H, Theiler J, Abfalterer W, Foley B, Giorgi EE, Bhattacharya T, Parker MD, Partridge DG. Spike mutation pipeline reveals the emergence of a more transmissible form of SARS-CoV-2. bioRxiv 069054v2 (Preprints) 2020. [Cited June 30, 2020]. Available from: https://www.biorxiv.org/content/10.1101/2020.04.29.069054v2.full

11. Hu J, He CL, Gao Q, Zhang GJ, Cao XX, Long QX, Deng HJ, Huang LY, Chen J, Wang K, Tang N. The D614G mutation of SARS-CoV-2 spike protein enhances viral infectivity and decreases neutralization sensitivity to individual convalescent sera. bioRxiv 161323v1 (Preprints) 2020. [Cited June 30, 2020]. Available from: https://www.biorxiv.org/content/10.1101/2020.06.20.161323v1.full

12. Wang Q, Zhang Y, Wu L, et al. Structural and Functional Basis of SARS-CoV-2 Entry by Using Human ACE2. Cell. 2020;181(4):894–904.e9.

13. Kirchdoerfer RN, Ward AB. Structure of the SARS-CoV nsp12 polymerase bound to nsp7 and nsp8 co-factors. Nat. Commun. 2019 May 28;10(1):1–9.

14. Yu IM, Oldham ML, Zhang J, Chen J. Crystal structure of the severe acute respiratory syndrome (SARS) coronavirus nucleocapsid protein dimerization domain reveals evolutionary linkage between corona-and arteriviridae. J. Biol. Chem. 2006 Jun 23;281(25):17134–9.

15. Zhao P, Cao J, Zhao LJ, Qin ZL, Ke JS, Pan W, Ren H, Yu JG, Qi ZT. Immune responses against SARS-coronavirus nucleocapsid protein induced by DNA vaccine. Virology. 2005 Jan 5;331(1):128–35.

16. Sanjuán R, Domingo-Calap P. Mechanisms of viral mutation. Cell Mol Life Sci. 2016 Dec 1;73(23):4433–48.

17. Di Giorgio S, Martignano F, Torcia MG, Mattiuz G, Conticello SG. Evidence for host-dependent RNA editing in the transcriptome of SARS-CoV-2. Sci. Adv. 2020 May 18:eabb5813.

18. Rosenberg BR, Hamilton CE, Mwangi MM, Dewell S, Papavasiliou FN. Transcriptome-wide sequencing reveals numerous APOBEC1 mRNA-editing targets in transcript 3’ UTRs. Nat Struct Mol Biol 2011 Feb;18(2):230.

19. Gonzalez-Reiche AS, Hernandez MM, Sullivan MJ, Ciferri B, Alshammary H, Obla A, Fabre S, Kleiner G, Polanco J, Khan Z, Alburquerque B. Introductions and early spread of SARS-CoV-2 in the New York City area. Science. 2020 May 29.

20. Regoes RR, Hamblin S, Tanaka MM. Viral mutation rates: modelling the roles of within-host viral dynamics and the trade-off between replication fidelity and speed. P Roy **Soc** B-**Biol** Sci 2013 Jan 7;280(1750):20122047.

21. Fitzsimmons WJ, Woods RJ, McCrone JT, Woodman A, Arnold JJ, Yennawar M, Evans R, Cameron CE, Lauring AS. A speed-fidelity trade-off determines the mutation rate and virulence of an RNA virus. PLoS Biol. 2018 Jun 28;16(6):e2006459.

22. Duffy S. Why are RNA virus mutation rates so damn high?. PLoS Biol. 2018 Aug 13;16(8):e3000003.

23. Sanjuán R, Thoulouze MI. Why viruses sometimes disperse in groups. Virus Evol. 2019 Jan;5(1):vez014.

24. Combe M, Sanjuan R. Variation in RNA virus mutation rates across host cells. PLoS Pathog. 2014 Jan 23;10(1):e1003855.

25. Zanini F, Neher RA. Quantifying selection against synonymous mutations in HIV-1 env evolution. J Virol. 2013 Nov 1;87(21):11843–50.

26. Stern A, Bianco S, Te Yeh M, Wright C, Butcher K, Tang C, Nielsen R, Andino R. Costs and benefits of mutational robustness in RNA viruses. Cell Rep. 2014 Aug 21;8(4):1026–36.

27. Silander OK, Tenaillon O, Chao L. Understanding the evolutionary fate of finite populations: the dynamics of mutational effects. PLoS Biol. 2007 Apr 3;5(4):e94.

28. Peck KM, Lauring AS. Complexities of viral mutation rates. J Virol. 2018 Jul 15;92(14).

